# Chromosome-level genome assemblies of *Nicotiana attenuata* and *Nicotiana obtusifolia*

**DOI:** 10.1101/2025.11.10.687602

**Authors:** Abhisek Chakraborty, Shuqing Xu

## Abstract

*Nicotiana attenuata* and *Nicotiana obtusifolia* are two wild tobacco species from the Solanaceae plant family, which are rich in producing diverse specialized metabolites and have been established as ecological model systems for studying plant-insect interactions. The previously published *N. attenuata* and *N. obtusifolia* genomes were draft assemblies and had low assembly contiguity, limiting comparative genomic analysis. Here, we performed chromosome-level genome assemblies and annotations of these two species using long-read sequencing and Hi-C data. Both *N. attenuata* and *N. obutosifolia* genomes were anchored to 12 chromosomes, with an assembled genome size of 2.19 Gb and 1.3 Gb, respectively. We achieved high BUSCO completeness at the protein level, with 99.3% for *N. attenuata* and 97.9% for *N. obtusifolia*. The reference genomes of *N. attenuata* and *N. obtusifolia* presented in this study will advance the understanding of metabolic innovations in Solanaceae plants.

## BACKGROUND & SUMMARY

*Nicotiana attenuata* and *Nicotiana obtusifolia* are diploid (2n = 2x = 24) Solanaceae species, which have been developed as ecological model plants to study plant-environment interactions in nature^1–3^. In particular, *N. attenuata* has been extensively used to understand the function, evolution and diversification of plant specialized metabolites (PSM), such as nicotine, as a defensive response to herbivory and other ecological factors, using genomics, transcriptomics, and metabolomics^3–5^. Further, *N. attenuata* has been used to understand the balance between plant defence and protection of autotoxicity conferred by the toxic PSMs^6^. Similarly, *N. obtusifolia* is also used to study ecological interactions of plants with pathogens, herbivores, and pollinators in natural habitats, along with *N. attenuata* and others^3^. Previous omics studies have used *N. obtusifolia* to understand the genetic basis of metabolic innovation and diversification of plant specialized metabolites in the *Nicotiana* genus, which are produced as a defence response to environmental stress^7–9^.

With the significance of these species as ecological models, and the necessity of future genomics-guided multi-omics strategies to further understand the metabolic innovations at the level of both plant organ and developmental stage, high-quality genomes are essential for comparative analysis^10^. A large number of genomic, transcriptomic, and metabolomic datasets already exist for *N. attenuata*^11^. However, high-quality genome assemblies were still not available for both of these *Nicotiana* species. The draft assemblies of *N. attenuata* and *N. obtusifolia* had low assembly contiguity (N50 values of 524.5 Kb and 134.1 Kb, respectively) in the previously published study^9^.

Here, we report the first chromosome-scale whole-genome assemblies of *N. attenuata* (12 chromosomes, 2.07 Gb) and *N. obtusifolia* (12 chromosomes, 1.29 Gb) (**Figure 1, Table 1**). Repetitive regions contributed to 83.26% and 79.21% of the *N. attenuata* and *N. obtusifolia* genome assemblies, respectively. After soft-masking the genome assemblies, we identified 35,166 and 27,352 protein-coding genes in *N. attenuata* and *N. obtusifolia*, respectively, using the BRAKER3 pipeline^12^ (**Table 1**).

**Table 1.**
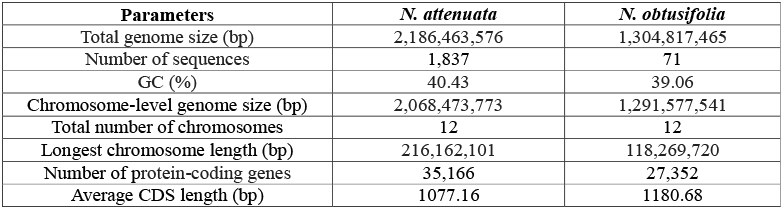
Genome assembly and annotation statistics of the *Nicotiana* genomes.

| Parameters | *N. attenuata* | *N. obtusifolia* |
| --- | --- | --- |
| Total genome size (bp) | 2,186,463,576 | 1,304,817,465 |
| Number of sequences | 1,837 | 71 |
| GC (%) | 40.43 | 39.06 |
| Chromosome-level genome size (bp) | 2,068,473,773 | 1,291,577,541 |
| Total number of chromosomes | 12 | 12 |
| Longest chromosome length (bp) | 216,162,101 | 118,269,720 |
| Number of protein-coding genes | 35,166 | 27,352 |
| Average CDS length (bp) | 1077.16 | 1180.68 |

**Figure 1.**
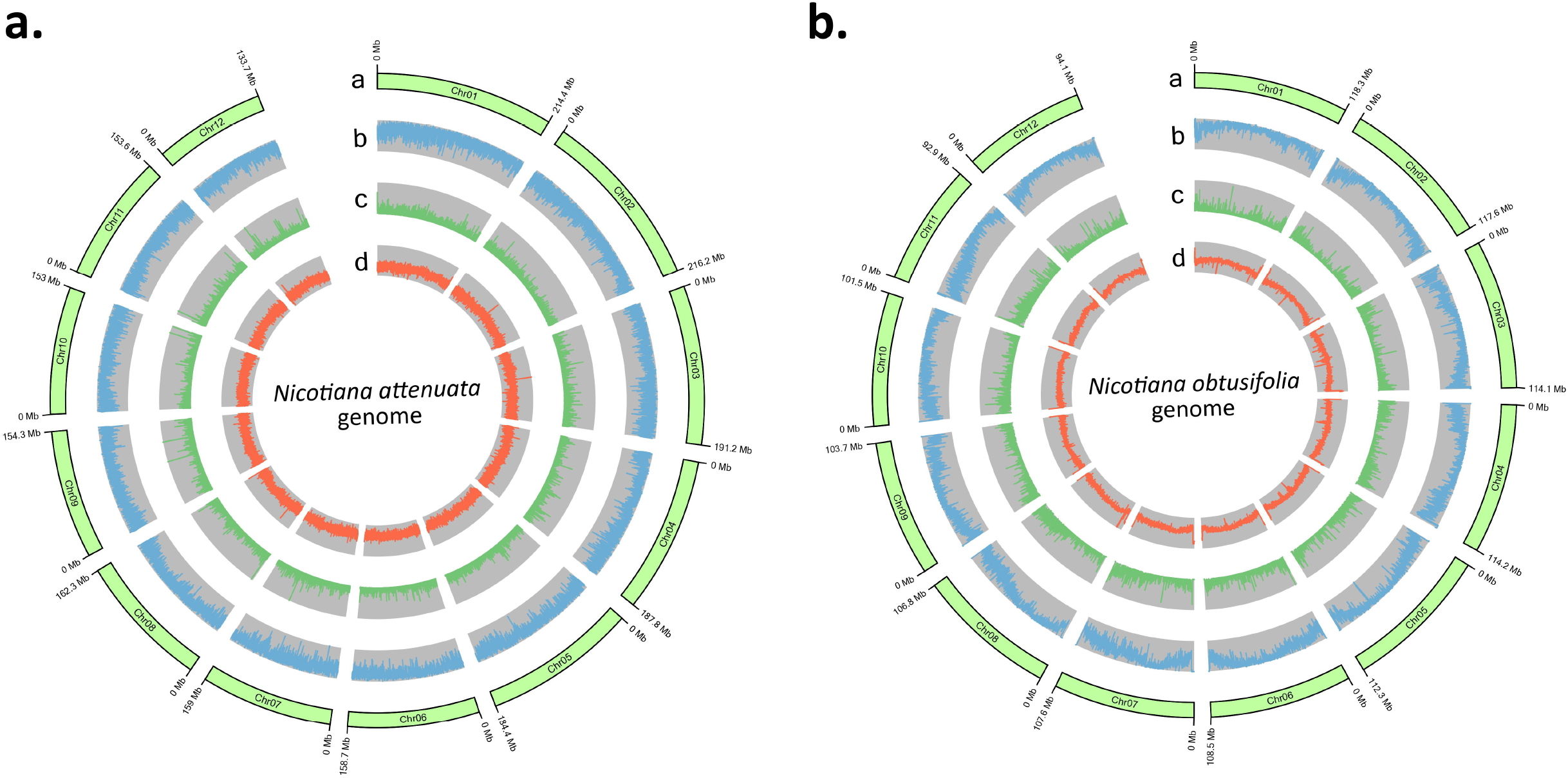
Circos plots showing the genomic features (window size 100 Kb)^45^. **A.** *N. attenuata* genome, **B**. *N. obtusifolia*. From the outer to inner circle – a. Chromosome length, b. Repeat density, c. Coding gene density, d. GC density.

We performed a whole-genome synteny analysis of *N. attenuata* and *N. obtusifolia* with previously published *N. sylvestris* (2n = 2x = 24) and *N. tomentosiformis* (2n = 2x = 24) genomes^13^, which showed notable genomic synteny among the *Nicotiana* species. However, chromosomal rearrangements were observed similar to other published *Nicotiana* genomes^14,15^, especially in the *N. obtusifolia* species (**Figure 2**).

**Figure 2.**
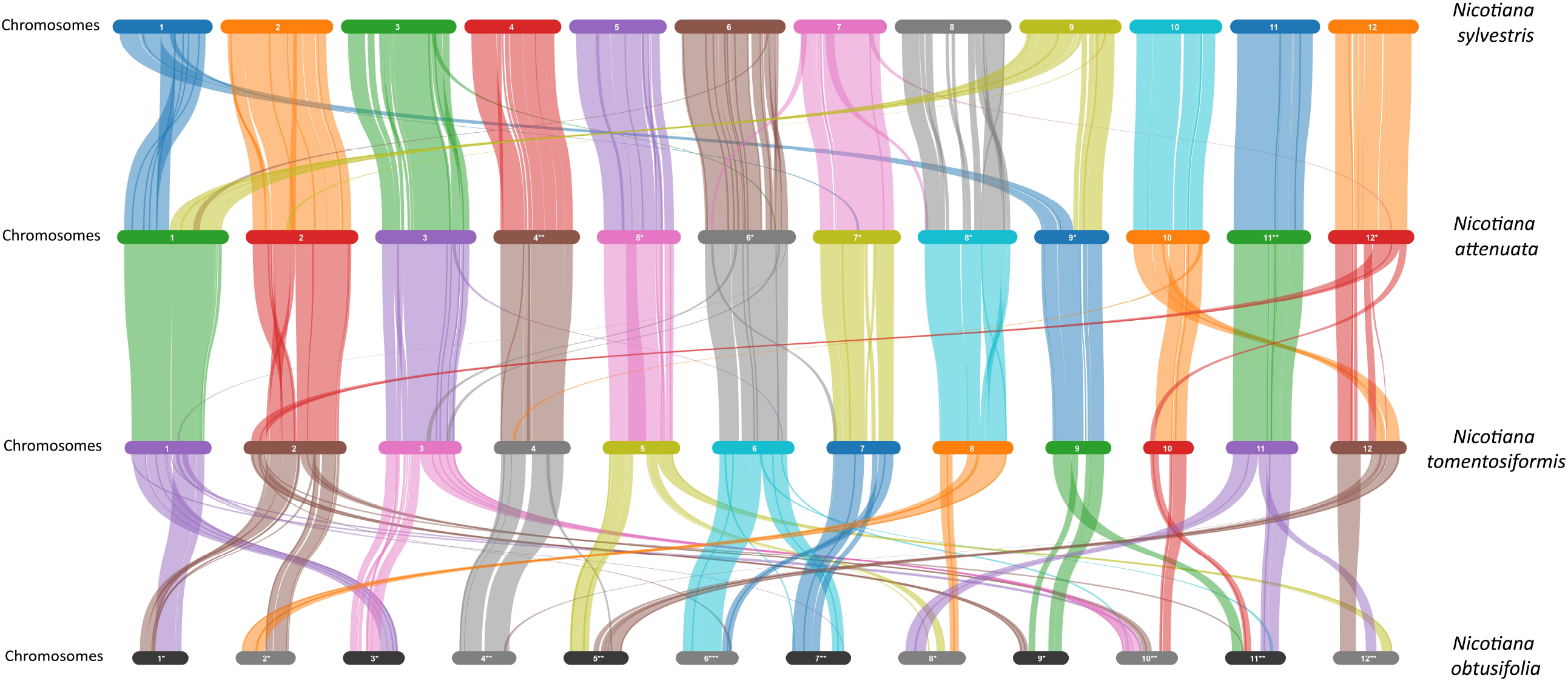
Genomic synteny analysis among four *Nicotiana* species – *N. attenuata* (this study), *N. obtusifolia* (this study), *N. sylvestris*^13^, and *N. tomentosiformis*^13^. The *N. attenuata* and *N. obtusifolia* chromosomes were renumbered and inverted relative to the *N. sylvestris* and *N. tomentosiformis* chromosomes for a better visualisation of the syntenic relationships. The single asterisks represent renumbered chromosomes, the double asterisks represent renumbered and inverted chromosomes, and the triple asterisk represents inverted chromosome.

The chromosome-level genome assemblies of *N. attenuata* and *N. obtusifolia* presented in this study will serve as a valuable dataset, facilitating a better understanding of the genomic basis of metabolic innovations not only in *Nicotiana* but also in the Solanaceae family.

## METHODS

### Sample preparation and sequencing

High-quality genomic DNA was extracted from frozen leaf samples using a modified CTAB method^16^. RNase A was used to remove RNA contaminants. For *N. attenuata*, a Hi-C fragment library (insert size 300-700 bp) was constructed for the whole genome and sequenced using the Illumina NovaSeq platform.

For *N. obtusifolia*, whole-genome sequencing was performed using the PacBio Revio platform, and for Hi-C-based contig anchoring, a Hi-C fragment library from 300-700 bp insert size was constructed and sequenced using the Illumina NovaSeq platform. PacBio Iso-seq RNA-seq library was also constructed using total RNA from the leaf sample, which was sequenced on a PacBio Sequel II platform. The sequencing was carried out at Biomarker Technologies (Beijing, China).

### Genome assembly

For *N. attenuata*, 301.92 Gb clean data were generated from the Hi-C library and were mapped to the previously assembled draft *N. attenuata* genome^9^ using BWA v0.7.17 with the default parameters^17^. Invalid read pairs, including Dangling-End and Self-cycle, Re-ligation and Dumped products, were filtered out. LACHESIS was used to cluster, order, and orient the contigs utilising the Hi-C interaction signals^18^. After Hi-C data-based anchoring, 2.07 Gb (94.6%) of the total assembly (2.19 Gb) was in confirmed order and orientation, constituting the 12 chromosomes. The parameters used in LACHESIS were: CLUSTER_MIN_RE_SITES = 161; CLUSTER_MAX_LINK_DENSITY = 2; ORDER_MIN_N_RES_IN_TRUNK = 199; ORDER_MIN_N_RES_IN_SHREDS = 239.

For *N. obtusifolia*, high-accuracy CCS data (96.91 Gb, N50 = 20.67 Kb) were assembled using Hifiasm v0.24.0 with the parameters: l=0 and n=4, resulting in 329 contigs^19^. For anchoring of the contigs, 197.74 Gb clean Hi-C data was mapped to the assembly using BWA v0.7.17 with default parameters^17^. Similar to the *N. attenuata* genome, the invalid read pairs were filtered out, and the valid interaction read pairs were used for the clustering, ordering, and orienting of scaffolds onto chromosomes with LACHESIS^18^. After Hi-C data-based anchoring, 1.29 Gb (98.99%) of the total assembly (1.3 Gb) was in confirmed order and orientation, constituting the 12 chromosomes. The parameters used in LACHESIS were: CLUSTER_MIN_RE_SITES = 723; CLUSTER_MAX_LINK_DENSITY = 2; ORDER_MIN_N_RES_IN_TRUNK = 15; ORDER_MIN_N_RES_IN_SHREDS = 15.

### Genome annotation

Both *N. attenuata* and *N. obtusifolia* genome assemblies were annotated using a similar method. Prior to coding gene prediction, the genome assemblies were used to construct the corresponding *de novo* repeat libraries using RepeatModeler v2.0.2, with “-LTRStruct” and other default parameters^20^. The resultant repeat libraries were used to soft-mask the respective genome assemblies using RepeatMasker v4.1.2 (https://www.repeatmasker.org).

The soft-masked *Nicotiana* genome assemblies were used for the prediction of high-confidence protein-coding genes using BRAKER3^12^, which implements an integrated transcriptome and proteome evidence-based gene prediction in GeneMark-ETP pipeline^21^. For transcriptome-based evidence of *N. attenuata*, the RNA-Seq data (Illumina short reads) from the previous study^9^ were quality-filtered using Trimmomatic v0.39 with the parameters “SLIDINGWINDOW:4:15 MINLEN:36”, and the filtered data were used in BRAKER3^12,22^. Inside the BRAKER3 pipeline, AUGUSTUS^23^ was also used for the prediction of coding genes, and TSEBRA^24^ was used for obtaining a combined output of GeneMark-ETP and AUGUSTUS-based gene predictions.

For transcriptome-based evidence of *N. obtusifolia*, both Illumina short read RNA-Seq data from a previous study^3^ and the newly generated PacBio Iso-seq ccs data (292,933 reads) were used in two separate BRAKER3 analyses. First, the Illumina short reads were quality-filtered using Trimmomatic v0.39 with the parameters “SLIDINGWINDOW:4:15 MINLEN:36”, and the filtered data were used in BRAKER3^12,22^. Next, the Iso-seq data were mapped onto the genome using Minimap2^25^, which was then used in a separate BRAKER3 analysis. The results from the two BRAKER3 analyses were merged using TSEBRA^24^.

In all the BRAKER3 analyses for both the *Nicotiana* species, the protein sequences from the previous draft assembly of *N. attenuata*^9^, along with species belonging to *Nicotiana* and other Solanaceae genus (only chromosome-level assemblies were considered) were used as a set of extrinsic proteome evidence in GeneMark-ETP. These Solanaceae species were - *N. sylvestris*^13^, *N. tomentosiformis*^13^, *N. tabacum*^13^, *N. benthamiana*^26^, *Capsicum annuum*^27^, *Solanum lycopersicum*^28^, *Physalis pruinosa*^29^, *Datura inoxia*^30^, *Lycianthes biflora*^30^, *Lycium chinense*^31^, and *Iochroma cyaneum*^32^.

The *Nicotiana* coding gene sets were filtered to extract the longest isoforms for each gene model and to remove the coding genes with a length of <100 bp using AGAT (https://github.com/NBISweden/AGAT). Further, the coding genes with a repeat content of >50% were filtered out, using a method similar to a previous study^33^. The protein-coding genes were annotated using the eggNOG-mapper genome annotation server, which uses orthology relationships to assign the KO (KEGG Orthology) terms, GO (Gene Ontology) terms, COG (Cluster of Orthologous Gene) categories, CAZy families, and Pfam domains^34^. Additionally, the coding genes were also mapped against the UniRef90 database using Diamond v2.0.13 with the parameters: “-k 1 -e 0.00001 -f 6 --sensitive”^35,36^.

### Inter-species genomic synteny analysis

Inter-species chromosomal synteny analysis was performed for the *N. attenuata* and *N. obtusifolia* genome assemblies from this study, with previously published chromosome-scale *N. sylvestris* and *N. tomentosiformis* genome assemblies^13^. MCScanX was used to analyse the inter-species collinearity among these four *Nicotiana* genomes with “-b 2” and “-s 15” parameters^37^, which was then visualised using SynVisio^38^.

### Evaluation of genome assembly and annotation completeness

To evaluate the completeness of the whole-genome assembly and the predicted coding gene sets, BUSCO v5.4.3 was used in “genome” and “proteins” modes, respectively, with the Solanales_odb10 database^39^. Secondly, the LTR Assembly Index (LAI) score^40^ was estimated for the chromosome-level genomes of *N. attenuata* and *N. obtusifolia* using GenomeTools^41^ v1.6.1 and LTR_retriever^42^ v3.0.1.

## DATA RECORDS

The newly generated Hi-C sequencing data for *N. attenuata* have been deposited in the NCBI database with BioProject accession number PRJNA1245670. Genome and transcriptome sequencing raw data of *N. obtusifolia* generated in this study have been deposited in the NCBI database with BioProject accession number PRJNA1332718. The genome assembly and annotation files are available at FigShare^43,44^. The mapping results of the *Nicotiana* gene sets constructed in this study against those of the previous study^9^ are also available at FigShare^43,44^.

## TECHNICAL VALIDATION

To evaluate the genome assembly quality and completeness, we implemented multiple strategies. First, the Hi-C contact map showed strong intra-chromosomal interaction signals along the diagonal, which confirms the genome structure integrity (**Figure 3**). Second, BUSCO analysis showed the presence of 98.5% complete BUSCO genes in both *N. attenuata* and *N. obtusifolia* genome assemblies, respectively (**Table 2**). At the proteome level, the BUSCO completeness was 99.3% and 97.9% for *N. attenuata* and *N. obtusifolia*, respectively (**Table 2**). Third, the chromosome-level genome assemblies of *N. attenuata* and *N. obtusifolia* had LAI scores of 14.6 and 15.6, respectively, which refer to “Reference”-standard genome assemblies, according to Ou et al.^40^ **(Figure 4**).

**Table 2.** BUSCO completeness of the *Nicotiana* genomes.

| Parameters | Genome | Proteins |
| --- | --- | --- |
| *N. attenuata* |  |  |
| Total BUSCO groups searched | 5,950 | 5,950 |
| Complete BUSCOs (C) | 5,861 (98.5%) | 5,909 (99.3%) |
| Complete and single-copy BUSCOs (S) | 5,648 (94.9%) | 5,725 (96.2%) |
| Complete and duplicated BUSCOs (D) | 213 (3.6%) | 184 (3.1%) |
| Fragmented BUSCOs (F) | 9 (0.2%) | 6 (0.1%) |
| Missing BUSCOs (M) | 80 (1.3%) | 35 (0.6%) |
| *N. obtusifolia* |  |  |
| Total BUSCO groups searched | 5,950 | 5,950 |
| Complete BUSCOs (C) | 5,861 (98.5%) | 5,829 (97.9%) |
| Complete and single-copy BUSCOs (S) | 5,646 (94.9%) | 5,630 (94.6%) |
| Complete and duplicated BUSCOs (D) | 215 (3.6%) | 199 (3.3%) |
| Fragmented BUSCOs (F) | 7 (0.1%) | 11 (0.2%) |
| Missing BUSCOs (M) | 82 (1.4%) | 110 (1.9%) |

**Figure 3.**
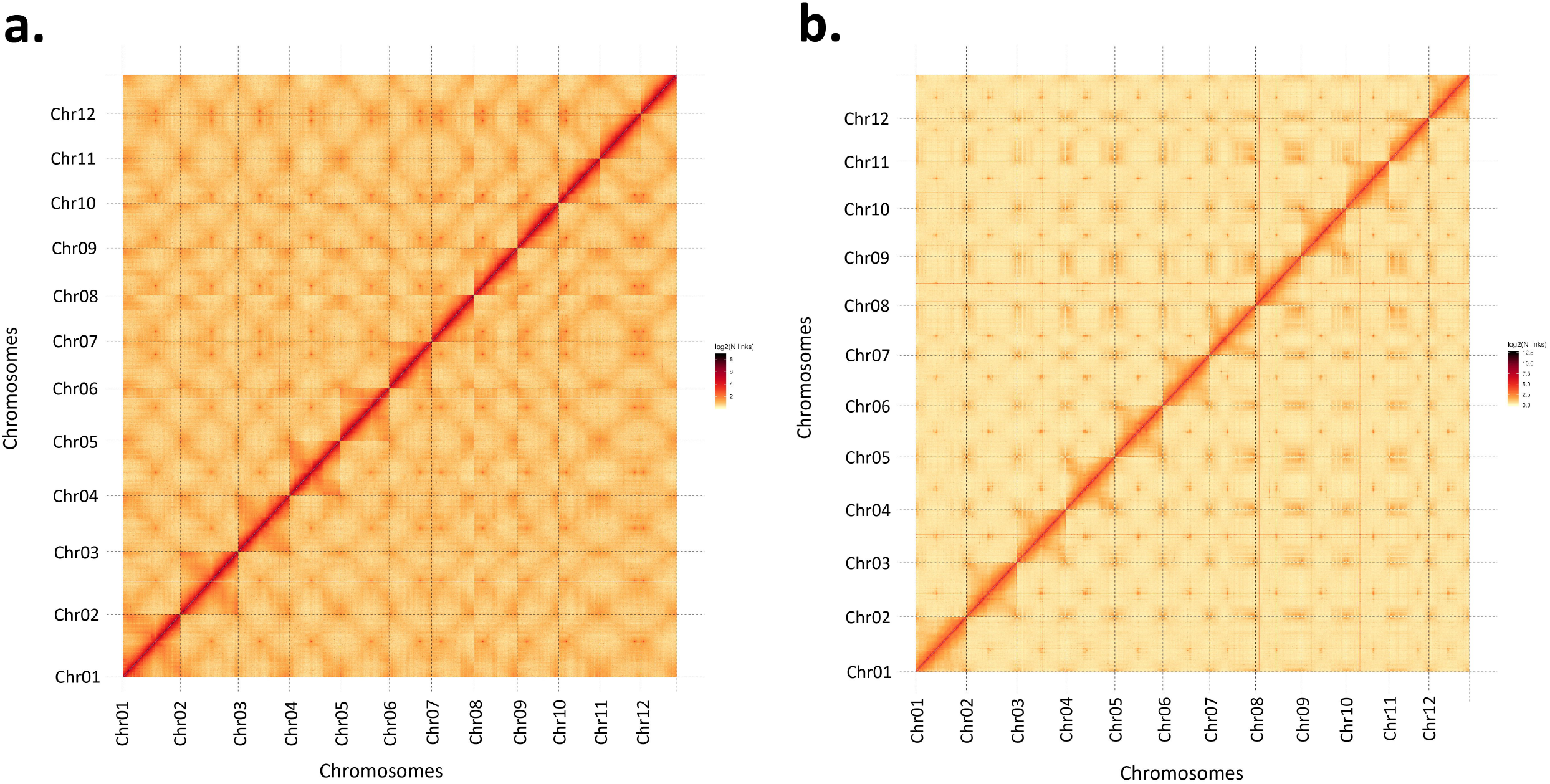
Hi-C contact maps of the genome assemblies - **A**.*N. attenuata* and **B**. *N. obtusifolia*.

**Figure 4.**
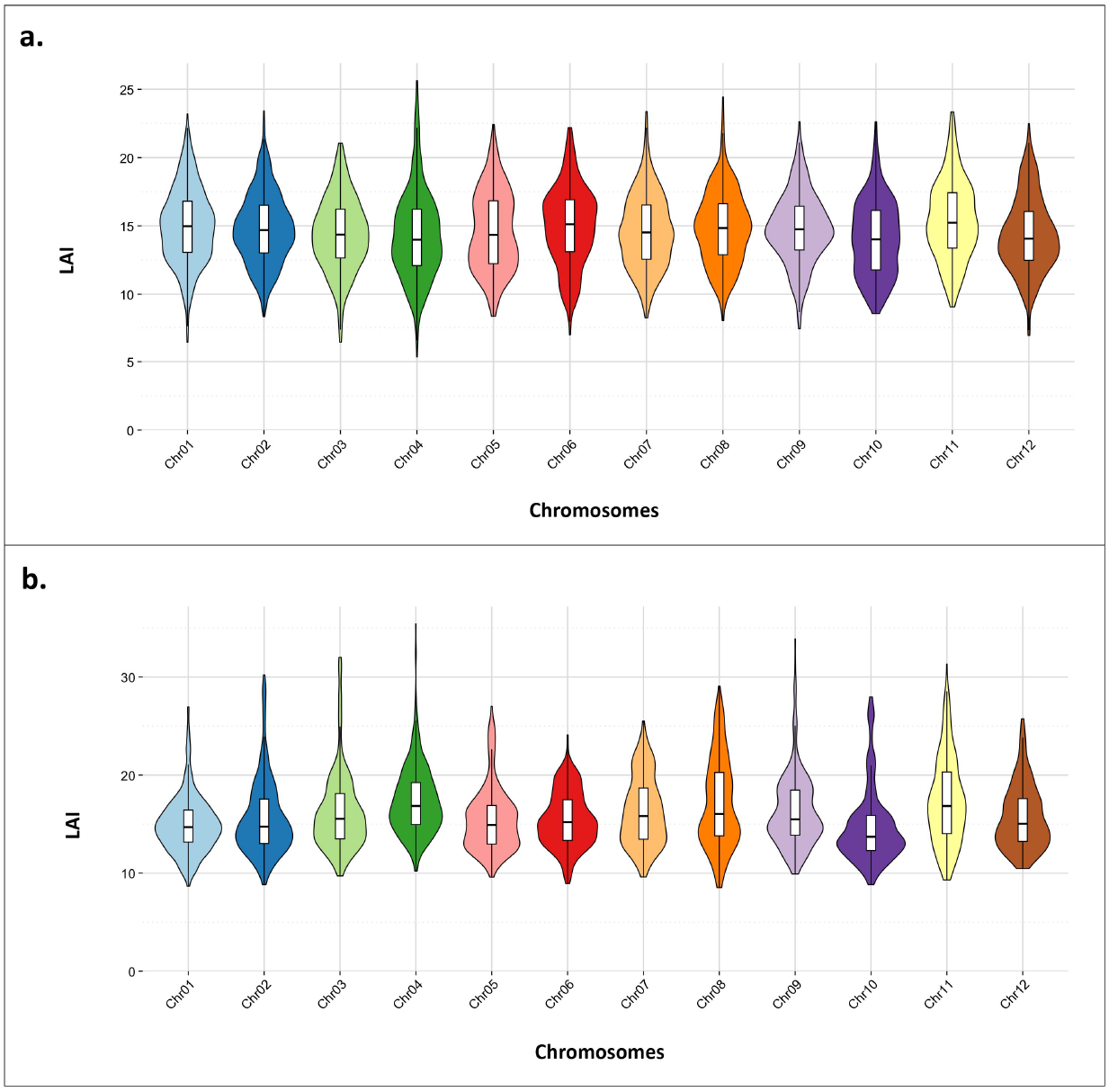
Distribution of the LAI scores in the 12 chromosomes of – **A.** *N. attenuata* and **B**. *N. obtusifolia*.

## CODE AVAILABILITY

All bioinformatic analyses were performed according to the manuals provided by the software developers. The software versions and the parameters used for the analyses are mentioned in the “Methods” section. The codes used in this study are deposited in GitHub with the link - https://github.com/Xu-lab-Evolution/Nicotiana_genome_project.

## ACKNOWLEDGEMENTS

We thank Dr. Danny Kessler from the Max Planck Institute for Chemical Ecology for providing the plant materials. We thank Dr. Patrycja Baraniecka and Thoomke Roth for preparing the plant materials for sequencing. Parts of this research were conducted using the supercomputer Mogon 2 and/or advisory services offered by Johannes Gutenberg University Mainz (hpc.uni-mainz.de), which is a member of the AHRP (Alliance for High-Performance Computing in Rhineland Palatinate, www.ahrp.info) and the Gauss Alliance eV. The authors gratefully acknowledge the computing time granted on the supercomputer MOGON 2 at Johannes Gutenberg University Mainz (hpc.uni-mainz.de).

## AUTHORS CONTRIBUTIONS

A.C. conducted the research and performed the data analysis. S.X. conceived and supervised the study. Both authors have read and approved the final manuscript.

## COMPETING INTERESTS

The authors declare no competing interests.

## FUNDING

This work was supported by the Agence Nationale de la Recherche (ANR)–Deutsche Forschungsgemeinschaft (DFG) joint research funding for French–German Collaboration (ANR-23-CE20-0037 to E.G. and DFG project number 529944545 to S.X.).

## Notes

### Competing Interest Statement

The authors have declared no competing interest.

